# In Vivo Non-Invasive Observation of Dynamic Changes in Hair Shaft, Melanin, and Collagen During the C57BL/6 Follicle Cycle

**DOI:** 10.1101/2025.09.15.676351

**Authors:** Menghua Liu, Gaiying He, Fenglong Wang, Yi Wang

**Affiliations:** Institute of Acupuncture and Moxibustion, China Academy of Chinese Medical Sciences, Beijing 100700, China; Experimental Research Center, China Academy of Chinese Medical Sciences, Beijing 100700, China; School of Life Science, Beijing University of Chinese Medicine, Beijing 102488, China

**Keywords:** Follicle, Second harmonic generation, anagen phase, C57BL/6 mice, alopecia

## Abstract

This study systematically monitored hair shaft, melanin, and collagen dynamics in vivo using non-invasive methods, enabling continuous observation over multiple hair follicle cycles. The hair shafts of C57BL/6 mice were observed in vivo using dermoscopy, photography, and electron microscopy. Second harmonic generation (SHG) and photography were employed for in vivo observation and quantification of melanin dynamics throughout the hair follicle cycle—specifically during the anagen phase, catagen phase, and telogen phase stages. Accuracy was validated via hematoxylin and eosin (H&E) staining. Additionally, SHG was used for in vivo observation and quantification of collagen changes across the same hair cycle phases, with validation performed using a liquid crystal polarizing imaging system and Sirius Red staining. This project provides non-invasive and in vivo methods for evaluating hair shafts, melanin, and collagen fibers across different growth cycles, thereby supporting non-invasive research in the development of active agents for hair loss treatment.

## Introduction

The hair follicles of C57BL/6 mice exhibit synchronicity and regularity, offering experimental convenience and making them a common animal model for studying skin diseases1,2.

The hair follicles enter the second telogen phase in the seventh week after the birth of the C57BL6 mouse^3^. In the absence of artificial intervention, this telogen phase will be maintained until the twelfth week after birth^4^. This extended period of telogen is a common time point used in research to synchronize the hair follicle growth cycle^5^. After artificial depilation of the mice, the hair follicles immediately begin to enter the anagen phase^6^.

After entering the growth phase, stem cells in the bulge region proliferate and differentiate into the outer root sheath, growing downward to increase follicle length^7^. The hair shafts base extends upward as the follicle deepens^8^. Stem cells in the bulb continuously differentiate into the hair shaft and inner root sheath^9^.The secretion of collagen and melanin in the skin also changes rapidly during the anagen phase^10^, playing an important role in hair follicle development^11,12^. In hair-loss-related studies, melanin secretion is generally assessed via skin color, which does not allow precise quantification^13^. So far, no consensus has been reached on how collagen content fluctuates across the follicular cycle.

Traditional methods for observing hair follicle cycles have significant limitations. Histological section staining requires euthanizing animals^14^. Conventional non-invasive techniques, such as dermatoscopy, are limited in depth and cannot visualize deep follicle changes^15^. Additionally, during late regression and resting phases, the melanin-deficient root of club hair becomes difficult to detect^16,17^. Liquid crystal polarimetry can detect changes in the direction and amount of collagen fibers around hair follicles^18,19^. Advances in optical imaging technologies now enable non-invasive, longitudinal observation of follicular changes^20,21^. Two-photon and second harmonic generation imaging can detect changes in hair shafts, melanin, and collagen in vivo^22-24^. Skin confocal microscopy can observe pigmentation and hair follicle regeneration in vivo^25,26^. Optical coherence tomography can measure skin thickness in vivo^27-29^. Overall, non-invasive in vivo observation of hair follicles needs further research^30^.

## Animals and Experimental Methods

### Declaration of Animal Use

Male C57BL/6 mice, aged 6-7 weeks and weighing (20 ± 2) g, were provided by Beijing Vital River Laboratory Animal Technology Co., Ltd. The animal license number is SCXK (Jing) 2021-0006. The mice were housed in a 12-hour light/12-hour dark environment with free access to food and water. All animal experimental procedures were approved by the Institutional Animal Care and Use Committee of the Experimental Research Center of China Academy of Chinese Medical Sciences (Protocol Code: ERCCACMS11-2210-03). Hair growth status was examined every other day from 2 days to 26 days after depilation.

### Skin Color and Hair Photography

After depilation, the backs of the mice were photographed every other day until day 26. Starting from day 10, the mice were depilated before taking photos of the skin color.

### Liquid Crystal Polarized Imaging

Skin tissue samples were taken from the back every other day from day 2 to day 26. Paraffin sections were prepared and de-waxed, and photographs were taken under a polarized light microscope. The angle of the polarizer was kept consistent for each shot.

### Second Harmonic Generation (SHG) Imaging

After anesthesia, C57BL/6 mice were fixed on the stage of a two-photon microscope. A 25 × water immersion objective lens with a numerical aperture of 1.05 was used to perform z-axis scanning of the dermal tissue. The excitation wavelength was 950 nm, with a step size of 2 μm. The image acquisition resolution was 1,024 × 1,024 pixels, and the acquisition speed was 4 μm/pixel. Scanning started from the upper surface of the dermis, with a depth of 45 μm, to observe changes in collagen fiber fluorescence signals.

### Histopathological Observation

Skin tissue samples were taken from the back every other day from day 2 to day The skin tissue was placed on a hard card to prevent curling and immersed in a test tube containing 4% paraformaldehyde. The tissue was then dehydrated and embedded to prepare 6 μm thick paraffin sections. Before de-waxing, the sections were heated on a slide warmer for 120 minutes. The skin sections were stained with hematoxylin and eosin (H&E) and Sirius Red according to the instructions of the staining kits. The sections were then mounted with neutral gum. The sections were observed using an inverted microscope.

### Electron Microscopy

Newly grown hair shafts were randomly plucked from the hair growth area on the back of the mice in each group. Three mice were selected per group, and the plucked hair shafts were fixed. Scanning electron microscopy was used to observe the hair shafts.

### Statistical Methods

Image parameters were measured using ImageJ software (V1.8.0). All data were statistically analyzed using SPSS 17.0 software. Comparisons between two groups were performed using t-tests. Bar charts were created using Prism v6.0, and results are expressed as the mean ± standard deviation (x ± s). Differences were considered statistically significant at *P* < 0.05.

## Results

### Mouse Hair Observation

Photographic observations of mouse hair growth are shown in Figure 1A. Hair began to emerge through the skin surface on day 10, with maintaining development from day 12 to day 16 (P<0.05). Hair coverage plateaued from day 18 to day 26 (P>0.05) (Figure 1B). Skin microscopy enabled non-invasive in vivo observation of hair growth changes. On day 2, the skin was uniformly pink and hairless. By day 8,hair that is about to reach the epidermis were visible. On day 10, the hair shaft was seen emerging through the epidermis. The hair shaft thickened progressively from day 10 to day 16, with the maximum hair diameter observed on day 16. By day 26, the skin reverted to a pink appearance without visible hair shafts (Figure 1A).

**Figure 1.**
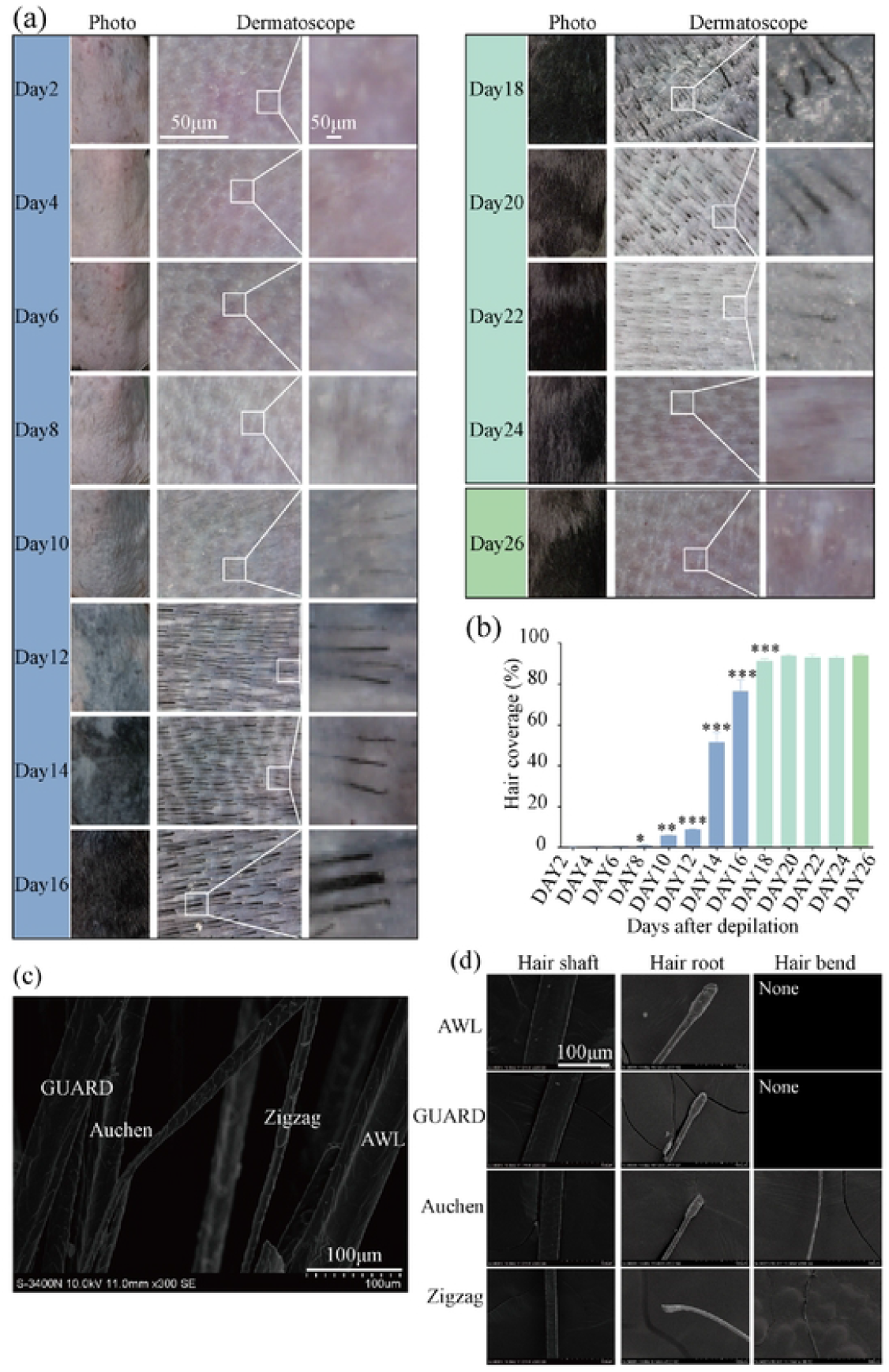
Observation of Hair Shafts in C57BL/6 Mice. (A) Photographic and dermatoscopic observation of mouse hair changes through the anagen (growth), catagen (regression), and telogen (resting) phases. (8) Statistical results of hair coverage observed by photography (n=3). (C) Scanning electron microscopy observation of four types of hair. (D) Characteristics of the four types of hair observed by scanning electron microscopy. Comparisons were made with the previous time points, with *P* <0.05, **P* <0.01, ***P* <0.001.

C57BL/6 mice develop four distinct hair types: GUARD, AWL, Auchen, and Zigzag, which are regularly distributed between larger primary and smaller follicles (Figure 1C)^31^. AWL and GUARD hairs are straight; AWL hairs are thicker and about half the length of GUARD hairs. Auchen hairs feature a single constriction. Zigzag hairs typically have two to four sharp bends, with segments arranged diagonally to form a wavy structure (Figure 1D).

### Observation of Melanin

Photographs were taken of the dorsal skin of the mice. The skin was pink on day 2, began to turn gray on day 6, turned black on day 12, started to gray again on day 18, and returned to pink by day 26. The experimental results are shown in Figure 2A. The gray value was statistically analyzed, and the optical density values of each day were compared with those of the previous day. A significant statistical difference was found starting from day 6 (P < 0.05). The skin gray value gradually increased from day 6 to day 16, indicating that the skin was gradually darkening. The gray value decreased from day 18 to day 22, indicating that the skin was returning to pink. The gray value remained unchanged on day 26 compared to the previous day, and the results are shown in Figure 2B.

**Figure 2.**
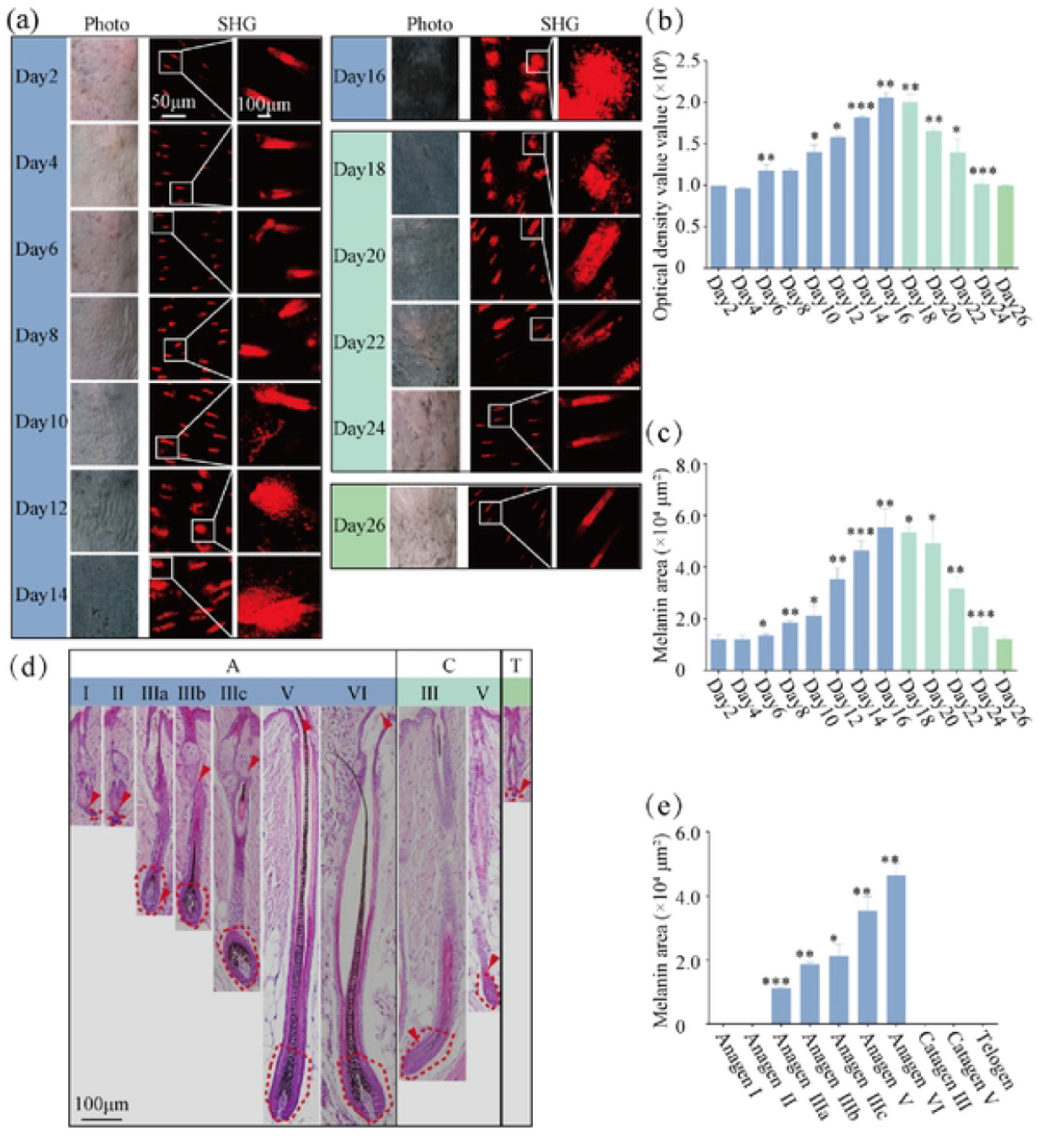
Observation of Melanin in C57BL/6 Mice. (A) Photographic and SHG observation of melanin changes through the anagen (growth), catagen (regression), and telogen (resting) phases. (8) Statistical results of skin gray value observed by photography (n=3). (C) Statistical results of melanin secretion observed by SHG (n=3). (D) HE staining observation of melanin in the hair follicle bulb. (E) Statistical results of melanin in the hair follicle bulb observed by HE staining (n=3). Comparisons were made with the previous time points, with *P* < 0.05, **P* <0.01, ***P* <0.001.

SHG in vivo observation technology was used to observe the changes in melanin secretion in the dorsal skin of mice, as shown in Figure 2C. Statistical analysis of the SHG results revealed that melanin secretion significantly increased on day 8. Melanin continued to increase from day 12 to day 16, reaching a maximum on day 16. It then gradually decreased starting from day 18, returning to the level of day 2 by day 24, as shown in Figure 2C.

H&E staining was used to observe melanin in the bulb of hair follicles at different growth stages, as shown in Figure 2D. Melanin secretion began at Anagen III and continuously increased throughout the growth phase. During the catagen and telogen phases, melanin in the hair follicle bulb completely disappeared, as shown in Figure 2E.

### Observation of Pores and Interpore Spaces

Collagen fibers in the interpore spaces were observed using Sirius Red staining and Abrio imaging (see Figure 3A). The results indicate that the collagen fibers between two pores began to become sparse on Day 6. The collagen continued to thin out from Day 8 to Day 16, reaching the minimum density on Day 16. Subsequently, the collagen density gradually recovered starting from Day 18 and returned to the Day 2 level by Day 24 (see Figures 3B-C).

**Figure 3.**
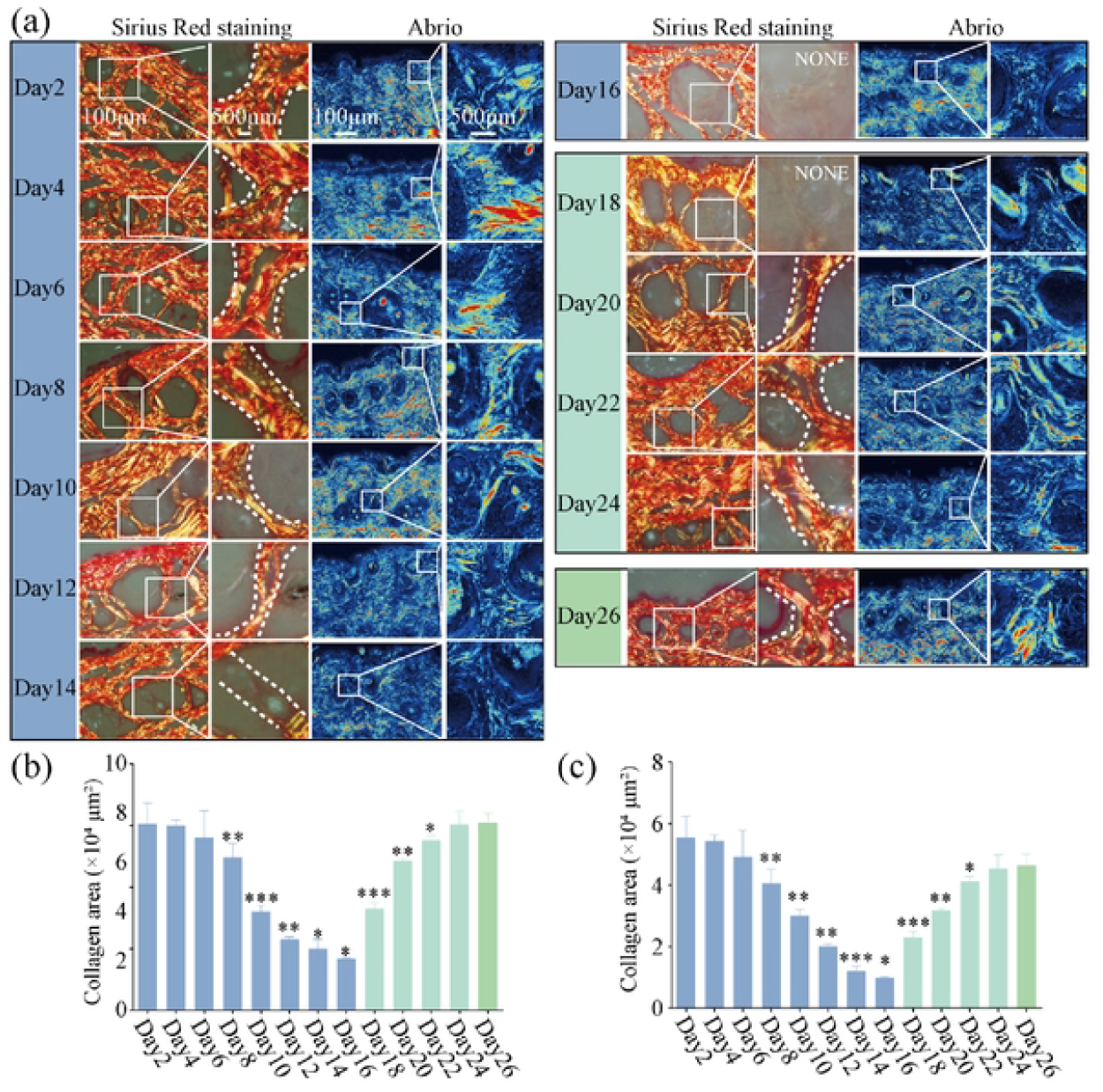
Histopathological Observation of Collagen Density in C57BL/6 Mice. (A) Sirius Red staining and liquid crystal polarized imaging observation of collagen changes in the interfollicular spaces through the anagen (growth}, catagen (regression), and telogen (resting) phases. (B) Statistical results of collagen density observed by Sirius Red staining (n=3). (C) Statistical results of collagen density observed by SHG imaging {n=3), Comparisons were made with the previous time points, with *P* < 0.05, **P* <0.01, ***P* <0.001.

SHG in vivo imaging was used to observe changes in collagen in the dorsal skin of mice, with observations made from the papillary layer, reticular layer, and Z-axis stacking (see Figure 4A). The papillary layer was observed at 5 μm, and the reticular layer at 23 μm (see Figure 4B). The collagen in the interfollicular spaces began to become noticeably sparse on Day 6. The density of collagen between pores continued to decrease from Day 8 to Day 16, reaching the minimum value on Day 16. Subsequently, it gradually recovered starting from Day 18 and returned to the Day 2 level by Day 24 (see Figure 4C).

**Figure 4.**
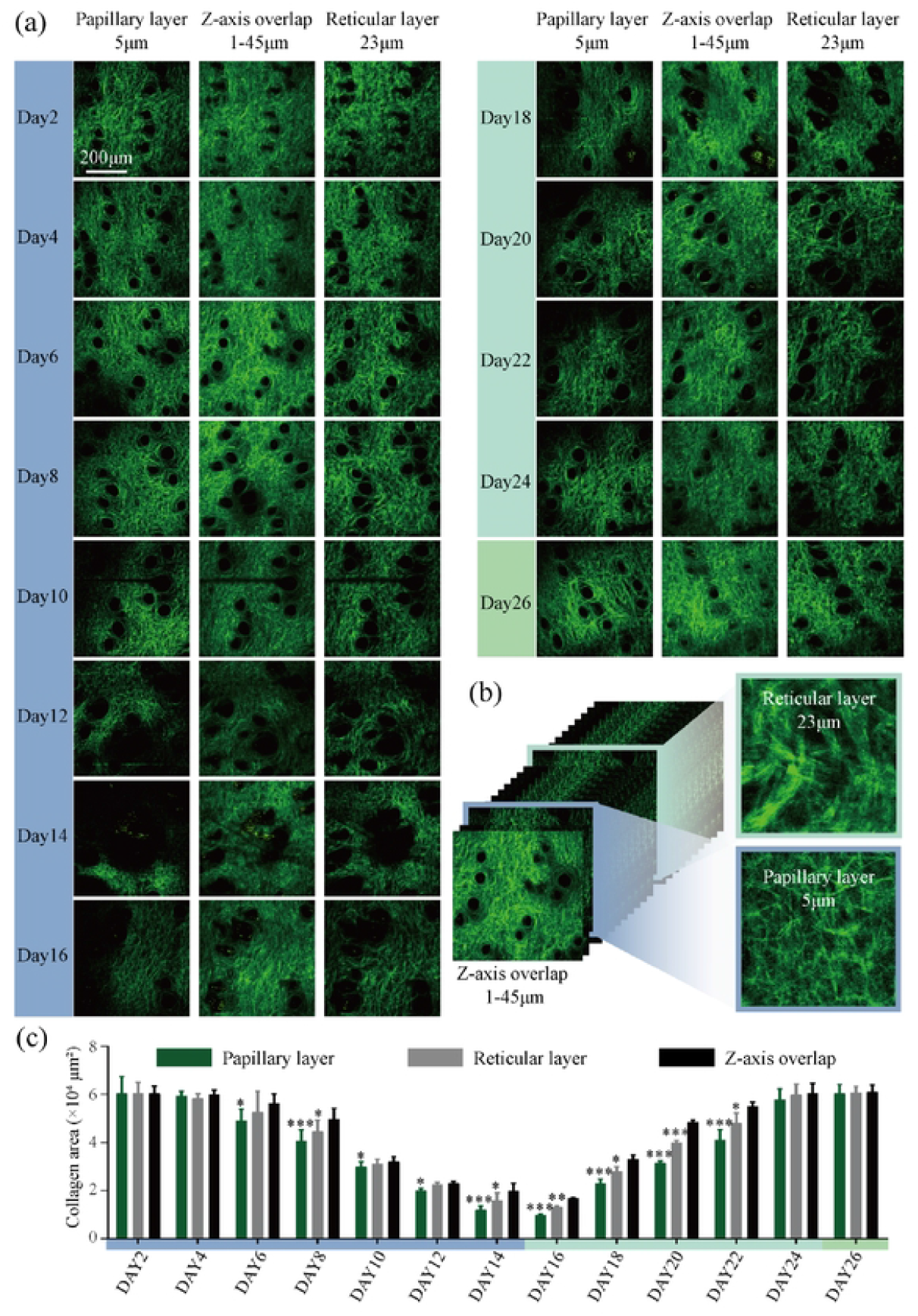
SHG In Vivo Observation of Collagen Density in C578L/6 Mice. (A) Observation of collagen changes in the interfollicular spaces through the anagen (growth), catagecn (regression), and telogen (resting) phases from the papillary layer, reticular layer, and Z-axis overlap. (8) Schematic representation of the selection and morphology of the papillary and reticular layers. (C) Statistical results of collagen density observed by SHG imaging (n=3). Comparisons were made with the previous time points, with *P* < 0.05, **P* <0.01, ***P* <0.001.

## Discussion

Regarding the hair follicle cycle in mice, when depilation is performed during the telogen phase, the hair follicles immediately enter the anagen phase. Innovatively, our research group was the first to establish a non-invasive in vivo method for observing the hair follicle cycle in mice based on two-photon and second harmonic generation imaging^32,33^. The hair follicles were found to be in the Anagen III stage on Day 6, in the Anagen VI stage on Day 10, entering the Catagen III stage on Day 18, the Catagen V stage on Day 20, and the telogen phase on Day 24, these results are consistent with those obtained from histopathological sections^4^. Moreover, since there is no need to sacrifice the mice, continuous multi-cycle studies can be conducted on the animals^34^. However, no systematic study has yet examined in vivo changes in melanin and collagen within the hair follicle growth cycle of C57BL/6 mice^34^. Building on our previous work, the present study therefore conducted longitudinal in vivo observations of these cyclical variations in melanin and collagen.

Regarding the study of hair shafts, the growth of hair was quantified by calculating the coverage rate of dorsal images through photography. The growth of hair was observed using dermatoscopy, with mouse hair beginning to emerge from the skin surface on Day 10 (Anagen VI stage). In previous research, the diameter of the hair shaft was quantified through non-invasive in vivo observation using two-photon microscopy. The diameter of the hair shaft at the skin surface continuously increased from Day 10 to Day 16 and gradually decreased during the catagen phase until it returned to the level of the telogen phase^33^.

Regarding the study of melanin, research has shown that the skin of depilated mice turns pink on Day 2, begins to gray on Day 6, turns black on the back on Day 12, starts to gray again on Day 18 (the beginning of the catagen phase), and turns pink on Day 26 (the telogen phase)^35^. This study used hematoxylin and HE staining to completely observe the expression of melanin in the hair bulb during the hair follicle cycle. Melanocytes that produce pigment do not exist within the epidermis of C57BL/6 mice, the pigmentation of their trunk is entirely derived from melanocytes in the hair follicles^36^. Therefore, the production of melanin in the hair bulb can represent the overall melanin production. In this study, we used SHG imaging to conduct non-invasive in vivo research on the changes in melanin during the hair follicle cycle in mice^37^. We quantified the expression of melanin in the skin by measuring the melanin area and validated the results using photographic methods.

The collagen above the isthmus of the hair follicle is closely associated with hair shaft emergence through the epidermis, but its dynamic changes during the anagen, catagen, and telogen cycles have not been quantitatively studied^38,39^. Literature reports mainly focus on the changes in collagen content in the skin of C57BL/6 mice with age, indicating that collagen content decreases with increasing age^40,41^. In this study, we used SHG imaging to non-invasively observe the collagen density within 45 μm below the epidermis between pores during the hair follicle cycle. We found that the collagen density between pores gradually decreased during the anagen phase and gradually returned to the telogen level during the catagen phase. Additionally, we discovered that the changes in collagen density in the papillary layer were more significant than those in the reticular layer, and the changes in the reticular layer were more pronounced than those in the Z-axis overlap. Therefore, the papillary layer is more suitable for observing changes in collagen fibers. Moreover, our research group previously conducted the first systematic study using two-photon microscopy and Abrio to statistically analyze pore area throughout the entire hair follicle cycle. The pores began to change before the hair shaft emerged. On Day 6, the pores were significantly enlarged, and the pore area continued to increase from Day 8 to Day 16, reaching a maximum on Day 16. Subsequently, the pore area began to gradually decrease on Day 18 and returned to the Day 2 level by Day 24^33^.

## Conclusion

This study non-invasively observed the changes in hair shafts, melanin, and collagen in C57BL/6 mice throughout the entire hair follicle cycle. During Anagen I-II, hair shafts had not yet begun to develop, and melanin production was undetectable. There were no significant changes in pore size or collagen density. During Anagen III-VI, hair shafts grew upward continuously until breaking through the epidermis. The diameter of the hair shaft at the epidermis gradually increased, melanin production progressively increased, pores gradually enlarged, and collagen became progressively sparse. During Catagen, hair shafts ceased development and moved toward the epidermis. The diameter of the hair shaft at the epidermis gradually decreased, melanin production was undetectable, pores gradually reduced in size, and collagen density gradually increased. During Telogen, some hair shafts were shed, melanin production was undetectable, and both pore area and collagen density returned to levels seen at the beginning of the anagen phase. In the future, by leveraging the advantage of non-invasive in-vivo imaging, we will conduct serial intravital observations of the hair follicles in fluorescent transgenic mice with labeled stem cells across multiple follicular cycles, laying the groundwork for further development of hair-loss therapeutics.

## Acknowledgments

This work was supported by Scientific and Technological Innovation Project of China Academy of Chinese Medical Sciences (NLTS2025004), the Fundamental Research Funds for the central public welfare reasearch institutes (JJPY2025002).

